# A window to the brain: ultrasound imaging of human neural activity through a permanent acoustic window

**DOI:** 10.1101/2023.06.14.544094

**Authors:** Claire Rabut, Sumner L. Norman, Whitney S. Griggs, Jonathan J. Russin, Kay Jann, Vasileios Christopoulos, Charles Liu, Richard A. Andersen, Mikhail G. Shapiro

## Abstract

Recording human brain activity is crucial for understanding normal and aberrant brain function. However, available recording methods are either highly invasive or have relatively low sensitivity. Functional ultrasound imaging (fUSI) is an emerging technique that offers sensitive, large-scale, high-resolution neural imaging. However, fUSI cannot be performed through adult human skull. Here, we use a polymeric skull replacement material to create an acoustic window allowing ultrasound to monitor brain activity in fully intact adult humans. We design the window through experiments in phantoms and rodents, then implement it in a participant undergoing reconstructive skull surgery. Subsequently, we demonstrate fully non-invasive mapping and decoding of cortical responses to finger movement, marking the first instance of high-resolution (200 μm) and large-scale (50 mmx38 mm) brain imaging through a permanent acoustic window.

## INTRODUCTION

Measuring brain function in adult humans is essential for neuroscience research and the diagnosis, monitoring, and treatment of neurological and psychiatric disease. However, existing brain recording techniques come with major trade-offs between sensitivity, coverage, invasiveness, and the ability to record from freely moving participants. Functional magnetic resonance imaging (fMRI) accesses the whole brain but has limited sensitivity (requiring averaging) and spatiotemporal resolution. It additionally requires the participant to lie in a confined space and minimize movements, restricting the tasks they can perform. Other non-invasive methods, such as scalp electroencephalography and functional near infrared spectroscopy, are affordable and portable. However, the signals they produce are limited by volume conduction or scattering effects, resulting in poor signal-to-noise ratios and limited ability to measure function in deep brain regions.

Magnetoencephalography has good spatiotemporal resolution but is limited to cortical signals. Intracranial electroencephalography and electrocorticography have good temporal resolution and better spatial resolution but are highly invasive. Intracranial electroencephalography requires electrodes inserted into the brain while electrocorticography requires implantation beneath the skull or dura. Implanted microelectrode arrays set the gold standard in sensitivity and precision by recording the activity of individual neurons and local field potentials. However, these devices are also highly invasive, requiring insertion into the brain. Moreover, they are difficult to scale across many brain regions and have a limited functional lifetime due to tissue reactions or breakdown of materials over time. To date, only severely impaired participants for whom the benefits outweigh the risk have used invasive recording technologies. There is a clear and distinct need for neurotechnologies that optimally balance the tradeoffs between invasiveness and performance (**Fig.1**).

**Figure 1.**
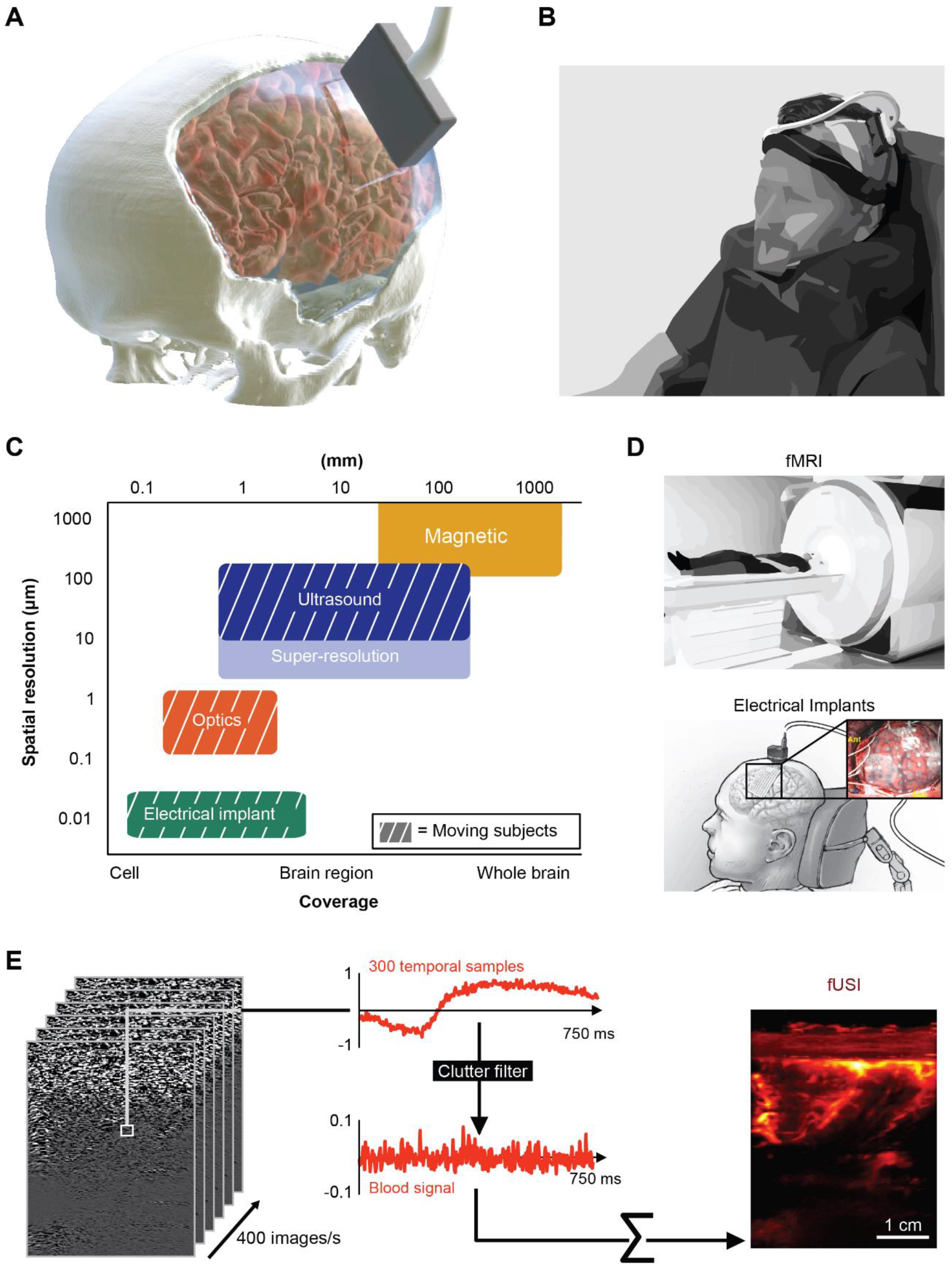
Permanent acoustic window enables noninvasive functional brain imaging in fully reconstructed, freely behaving subjects with high spatiotemporal resolution and large coverage. (**A**) Illustration of fUSI recording through a cranial window (**B**) Experimental setup for functional recording in a human using fUSI (**C**) Common functional recording modalities on a chart comparing their spatial resolution, coverage, and ability to record moving subjects (**D**) Example experimental setups for recording in humans using fMRI or electrical recording implants. ECoG sketch modified from ECoG sketch created by Ken Probst/UCSF (**E**) The ultrafast acquisition of ultrasound images allows a fast temporal sampling of the brain signal. We then applied a clutter filter to exclude tissue motion while keeping blood motion for sensitive measurements of CBV variations. Right panel: 2D rendering of fUSI recording through the participant’s scalp.

Functional ultrasound imaging (fUSI) is an emerging neuroimaging technique that spans the gap between invasive and non-invasive methods (**Fig. 1A-D**). It represents a new platform with high sensitivity and extensive brain coverage, enabling a range of new pre-clinical and clinical applications. Based on Power Doppler imaging, fUSI measures changes in cerebral blood volume (CBV) by detecting the backscattered echoes from red blood cells moving within its field of view (several cm) (**Fig. 1E**). It is spatially precise down to ∼100 μm with a framerate of up to 10 Hz, allowing it to sense the function of small populations of neurons^1^. fUSI is minimally invasive and requires only removal or replacement of a small skull area in large organisms. fUSI does not intrude on brain tissue but instead sits outside the brain’s protective dura mater and does not require the use of contrast agents. fUSI is non-radiative, portable, and proven across multiple animal models (rodents, ferrets, birds, non-human primates, and humans)^2,3^. In recent work, we decoded the intentions and goals of non-human primates from fUSI data^4^ and subsequently used fUSI as the basis for the first ultrasonic brain-machine interface (BMI)^5^.

An important direction of this research is the translation of fUSI-based neuroimaging and BMI for human participants.

However, the skull bone attenuates and aberrates acoustic waves at high frequencies, substantially reducing signal sensitivity. As a result, most pre-clinical applications require a craniotomy^5^, and the few human fUSI studies have required the skull to be removed or absent. These include intra-operative imaging during neurosurgery^7,8^ and recording through the anterior fontanelle window of newborns^9^ . Using fUSI to record brain activity in awake adults outside of an operating room is currently impossible.

In this study, we demonstrate fUSI in an awake adult participant equipped with an ultrasound-transparent “acoustic window” installed as part of a skull replacement procedure following a decompressive hemicraniectomy (partial skull removal) (**Fig. 1A**,**B**). Hemicraniectomies are commonly performed to reduce pathologically high intracranial pressures, including from traumatic brain injuries (TBIs), strokes, and subarachnoid hemorrhages^10–12^. After the craniectomy, the patient is often discharged from the hospital to home, rehabilitation, or care facilities with the skull defect covered by scalp for several weeks or longer depending on their clinical progress. After this period, a cranioplasty is performed to replace the missing skull with one of an assortment of reconstruction materials. These include autologous bone, titanium mesh, and polymethyl methacrylate (PMMA). Recently, customized cranial implants (CCIs) have grown in popularity thanks to their sterility, strength, and cosmetic appeal^13^. One CCI material, PMMA, is also purported to be transparent to ultrasound^14^, or “sonolucent”. This prompted us to ask whether we could design a skull replacement window to perform fUSI non-invasively in fully reconstructed adult humans, providing a convenient method to monitor brain health and giving access to human neural activity outside the operating room for neuroscience research and the development of brain-machine interfaces.

To test this possibility, we first examined the suitability of two FDA-approved skull replacement materials (PMMA and titanium mesh) for functional ultrasound imaging an in vitro cerebrovascular phantom, then compared their signal and contrast properties in an in vivo rodent model. This allowed us to design a PMMA acoustic window that could be permanently installed in a human patient as part of a skull reconstruction. Through this window and overlaying intact scalp, in an ambulatory setting outside the operating room, we demonstrated fully noninvasive recording and decoding of functional brain signals while our human participant performed visuomotor tasks, including playing a video game and strumming a guitar.

## RESULTS

### PMMA allows thickness-dependent blood flow imaging

fUSI is performed by acquiring a series of sequential power Doppler images and observing spatiotemporal changes in signal. To determine if fUSI signals can be detected through PMMA material, we first constructed a Doppler ultrasound phantom with flow channels designed to mimic blood flow in a human brain (**Fig. 2A**). This allowed us to measure the signals underlying fUSI in a controlled environment. We compared five different imaging scenarios: (1) no implant, (2) 1 mm thick PMMA implant, (3) 2 mm thick PMMA implant, (4) 3 mm thick PMMA implant, and (5) titanium mesh implant (**Fig. 2B-C**). We passed synthetic red blood cells through a 280-μm diameter tubing at three lateral (5, 15, 25 mm) and four axial positions (14, 24, 34, 44 mm) at a constant velocity of ∼27 mm/s, imaged them with a linear ultrasound array transmitting at 7.5 MHz, and recorded power Doppler signals to estimate the signal-to-noise ratio (SNR) and resolution loss in each imaging scenario (**Fig. 2D**). We found that signal intensity decreased with increasing PMMA implant thickness, and that image quality was most strongly degraded by the titanium mesh (**Fig. 2D-E**). The SNR decreased with depth and was inversely proportional to the thickness of the intervening PMMA material (**Fig. 2F**).

**Figure 2.**
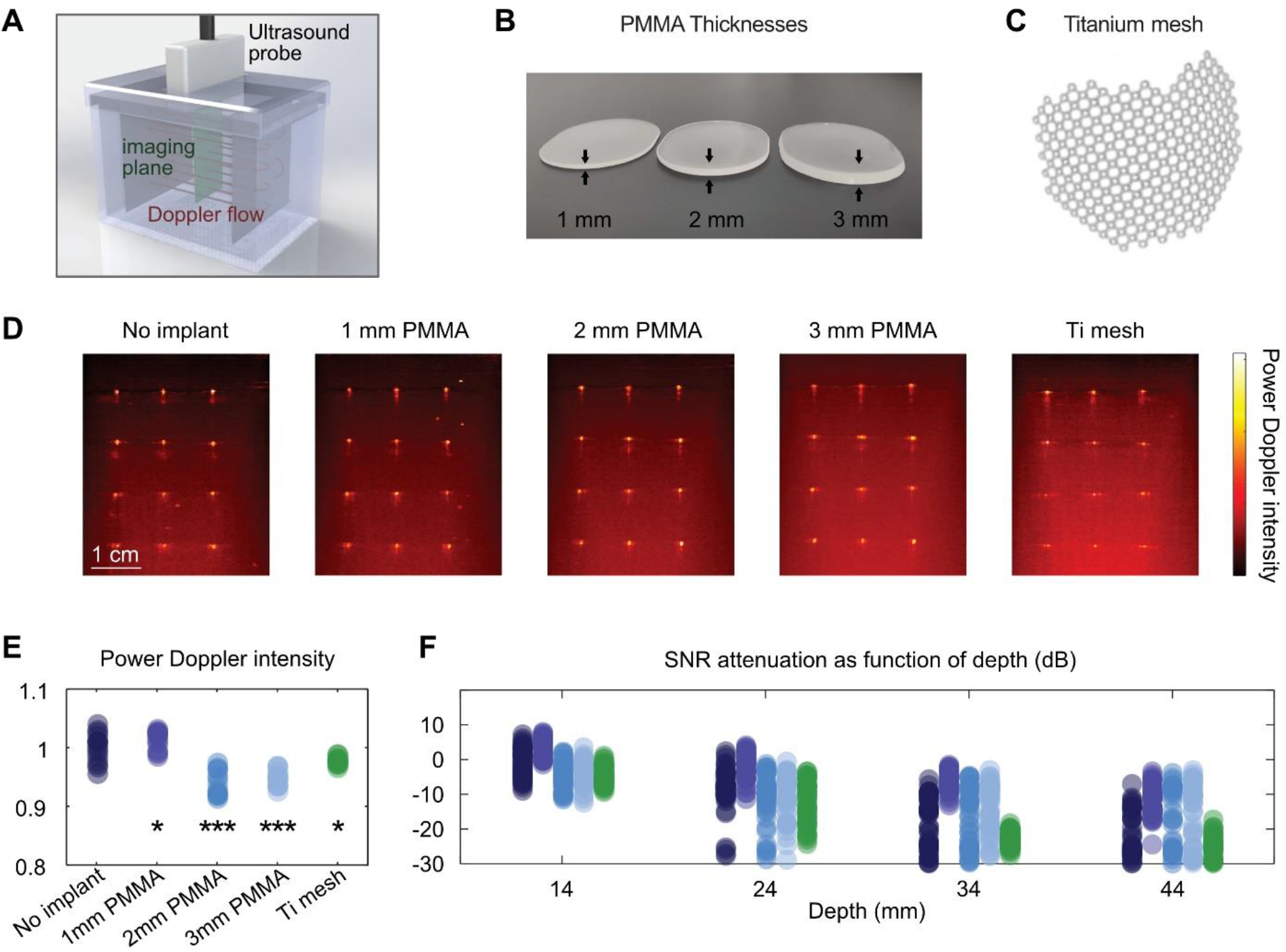
Polymeric skull replacement materials enable fUSI imaging with minimal SNR loss in an in vitro blood flow phantom. (**A**) 3D rendering of the in vitro Doppler phantom (**B**) Photo of three thicknesses of the PMAA skull-implant material used to evaluate the performance of the in vitro power Doppler recording through the implants (**C**) Photo of titanium mesh commonly used in cranial reconstruction (**D**) Power Doppler images (see A. for imaging plane) of the Doppler phantom in different implant scenarios (**E**) Power Doppler intensity for each implant scenario over N=15 acquisitions. Stars indicate statistical differences (paired sampled t-test) between each scenario compared to the No implant scenario (* : p < 0.05, **: p < 0.01, ***: p < 0.001) (**F**) SNR attenuation for each implant scenario as function of the depth (N=15 acquisitions).

### Skull replacement window allows fUSI imaging in a rodent model

To test our ability to detect functional brain signals through the different cranial implant materials in vivo, we performed fUSI in four rats after placing each of the five implant types on top of their brain following an acute craniotomy (**Fig. 3A**). We numerically evaluated the in vivo performance of fUSI through the implants by first calculating the overall fUSI signal intensity received through the different PMAA thicknesses and through the titanium mesh. The total fUSI intensity from the whole brain decreased by 30 % from the no-implant scenario to the 1 mm implant (**Fig. 3C**). The fUSI intensity dropped a further ∼15% per mm implant thickness for the 2 mm and 3 mm materials. The fUSI intensity decreased by 60 % for the titanium mesh compared to no implant. We then calculated the SNR of the cerebral blood vessels captured with the fUSI sequence (**Fig. S1**). **Figure 3C.ii-iii** shows that the SNR decreased as the PMMA implant thickness increased but with a smaller decrease than observed for fUSI intensity. In the cortex, the SNR decreased slightly with the mesh (-1 dB) and as the PMMA implant thickened (∼ -1 dB/mm). The subcortical structures within the image showed a similar trend in SNR across the different implant materials.

**Figure 3.**
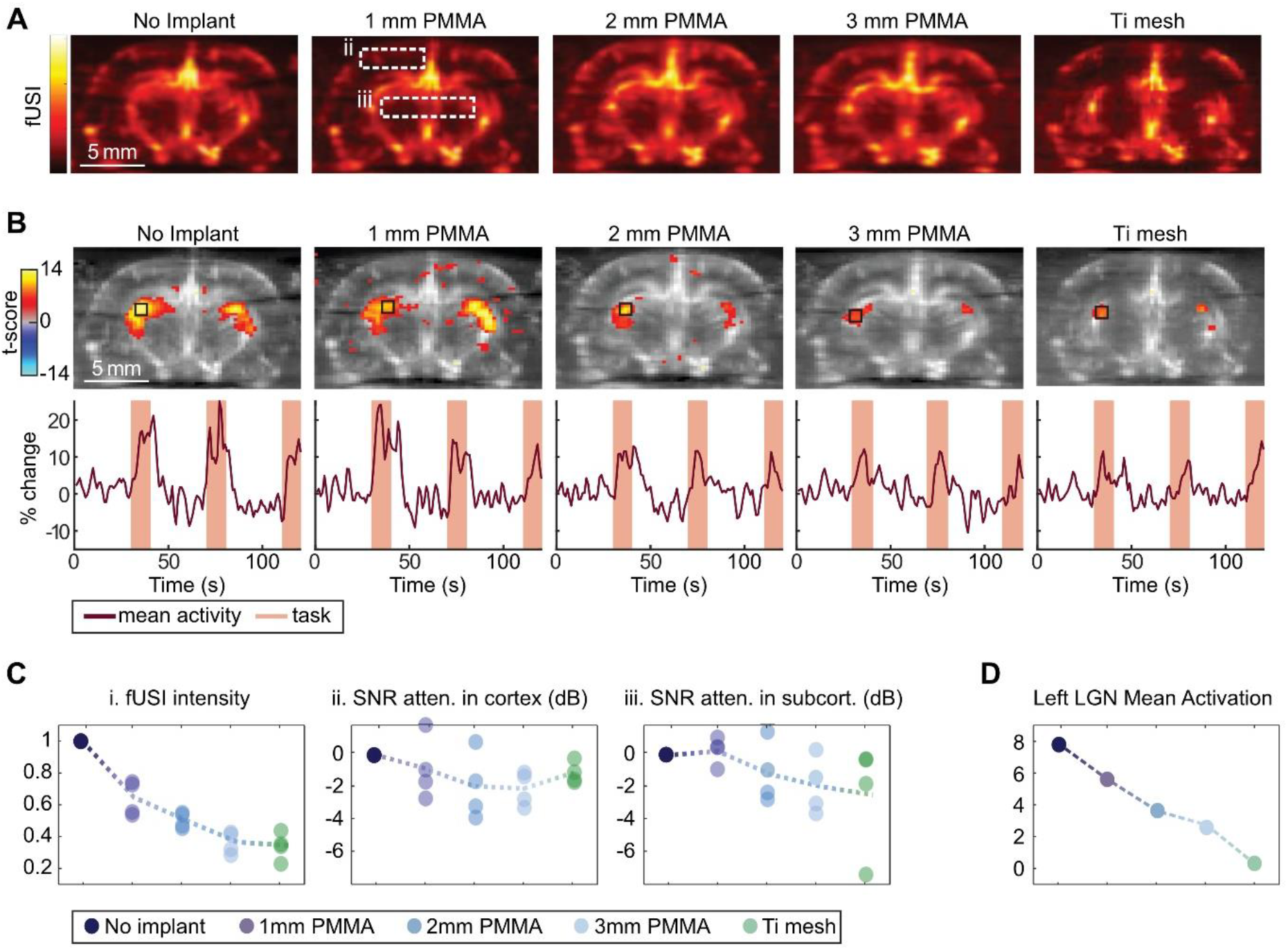
Thinner polymeric acoustic window materials enable the most sensitive in vivo rodent fUSI imaging. (**A**) fUSI images of the same rat brain with different skull implant materials (**B**) Areas modulated by visual stimulus, and example time courses. Top row - Statistical parametric map of voxels modulated by visual stimulation (p(corrected)<10-5) from GLM. Black box shows LGN region used to calculate mean fUSI time course. Bottom row - Time course from LGN region for each skull implant condition. Maroon line - Mean percent change. Orange shading - Light-on condition (**C**) Standardized fUSI intensity, standardized SNR in cortex (see dashed box ii. of panel A for cortex region), and standardized SNR in subcortical structures (see dashed box iii. of panel A for subcortical structures region) for each skull implant scenario (**D**) Significant activated area of the left LGN (in mm2) following visual stimulation as a function of the implant.

As we recorded the fUSI signals, we used a passive visual simulation task designed to activate the visual system. The rat experienced subsequent blocks of darkness (50-second) and light exposure (16.5-second). We modeled the response of each voxel to the visual stimulation using a general linear model (GLM), which allowed us to quantify which voxels showed significant visual modulation (p<10^-5^). Briefly, we convolved our block design (“rest” or “light”) with the hemodynamic response function and fit the linear model mapping of the convolved stimulation regressors to each fUSI voxel’s signal. This allowed us to assess the statistical significance between the hemodynamic response and the stimulation structure for each voxel. In all five implant conditions, we identified voxels within the lateral geniculate nucleus (LGN) activated during optical stimulation (**Fig. 3B**). Using fUSI through the thicker implants and the titanium mesh resulted in fewer activated voxels within the LGN. Additionally, we observed less signal change for the thicker vs. thinner implants and titanium mesh vs. any of the PMMA implants (p<10^-3^, 1-way ANOVA with post hoc hsd, **Fig. 3D**). Taken together, our in vitro and in vivo results suggested that PMMA was superior to the titanium mesh as an intervening material for fUSI and that making the PMMA window as thin as possible would offer the best imaging performance.

### Power Doppler images can be acquired through the scalp prior to skull reconstruction

To test the possibility of performing fUSI through a chronic cranial window, we recruited a human participant – an adult male in his thirties. Approximately 30 months prior to skull reconstruction, Participant J suffered a traumatic brain injury and underwent a left decompressive hemicraniectomy of approximately 16 cm length by 10 cm height (**Fig. 1A**). Anatomical and functional MRI scans allowed us to map brain structures and functional cortical regions within the borders of his craniectomy (**Fig. 4A**,**B**).

**Figure 4.**
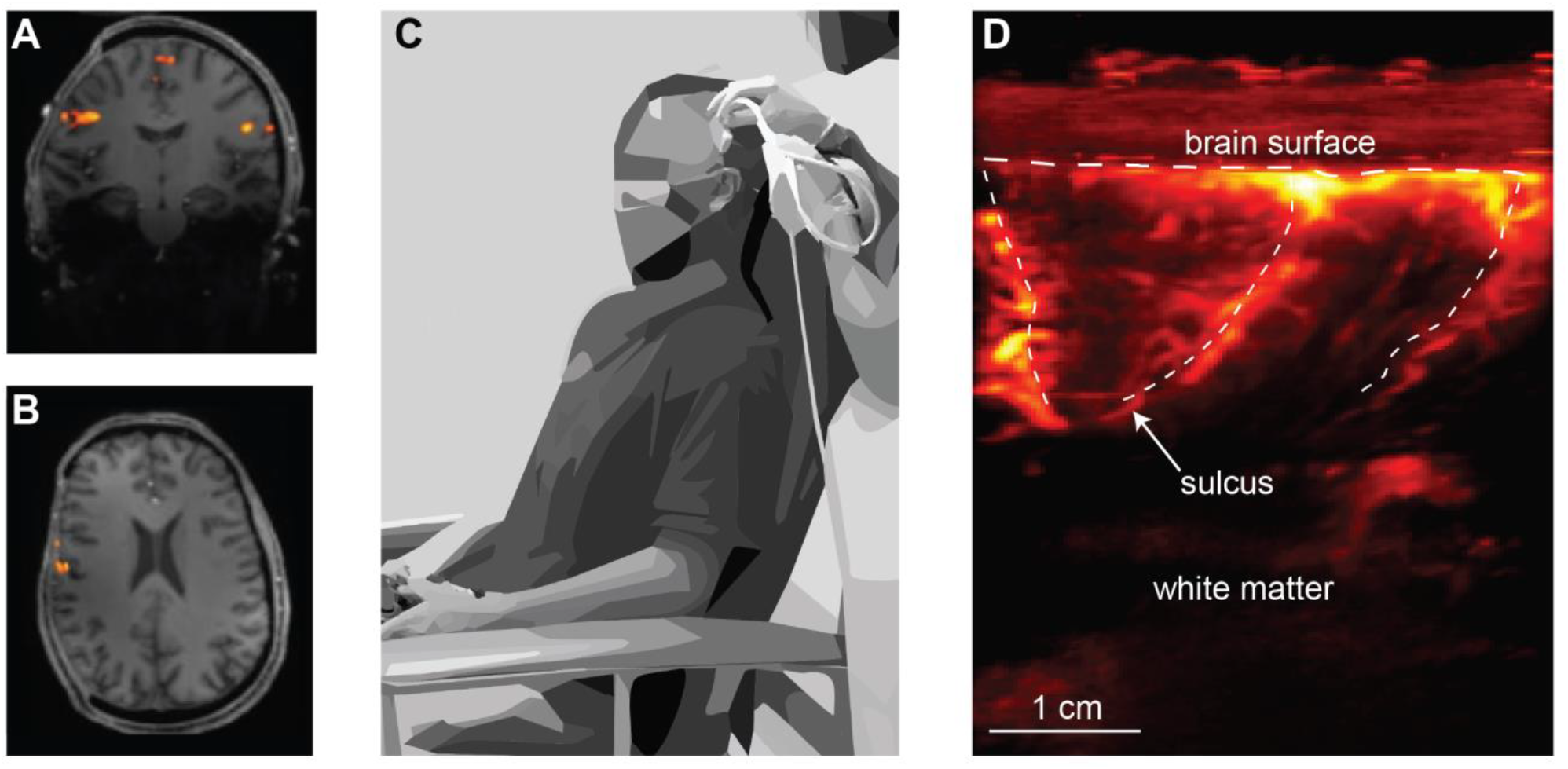
Ultrasound enables vascular imaging through intact scalp after decompressive craniectomy. (**A**) Coronal plane from Participant J’s fMRI after their decompressive hemicraniectomy. Orange overlay shows regions activated during finger tapping task (**B**) Transverse plane showing finger-tapping regions (**C**) Participant J during functional ultrasound imaging session with craniectomy (D) Power Doppler image of Participant J’s brain through the scalp.

Before reconstruction, we imaged participant J’s brain using power Doppler ultrasound through his intact scalp, with no intervening bone (**Fig. 4C**). We observed large brain vessels following the curve of sulci folds and smaller vessels irrigating the sulci, typical of fUSI images (**Fig. 4D**). However, due to the lack of intracranial pressure and the dramatic brain motion that results from this condition, we were unable to collect functional data or co-register ultrasound images to anatomical MRIs. Nevertheless, the ability to collect high quality vascular maps provided evidence that fUSI was possible through an intact human scalp and motivated us to proceed with designing, installing, and testing an acoustic window for Participant J.

### Customized cranial implant allows the installation of a 2 mm-thick PMMA window

To successfully detect functional signal through Participant J’s CCI, we collaborated with his attending physician (author CL) and the CCI manufacturer to design an appropriate acoustic window. In a separate fMRI study, we identified cortical response fields to a simple finger tapping task prior to his skull reconstruction (**Fig. 4A**,**B**). Based on this mapping, his attending physician and the CCI manufacturer manufactured the PMMA CCI implant with a 2 mm-thick 34 mm x 50 mm parallelogram-shaped sonolucent “window to the brain” (**Fig. 5A**,**B**). The 2 mm-thick portion sits above the primary motor cortex, primary somatosensory cortex, and posterior parietal cortex. The remaining PMMA implant is 4 mm-thick. This implant design was calculated by the manufacturer to provide sufficient mechanical performance to serve as a permanent skull replacement.

**Figure 5.**
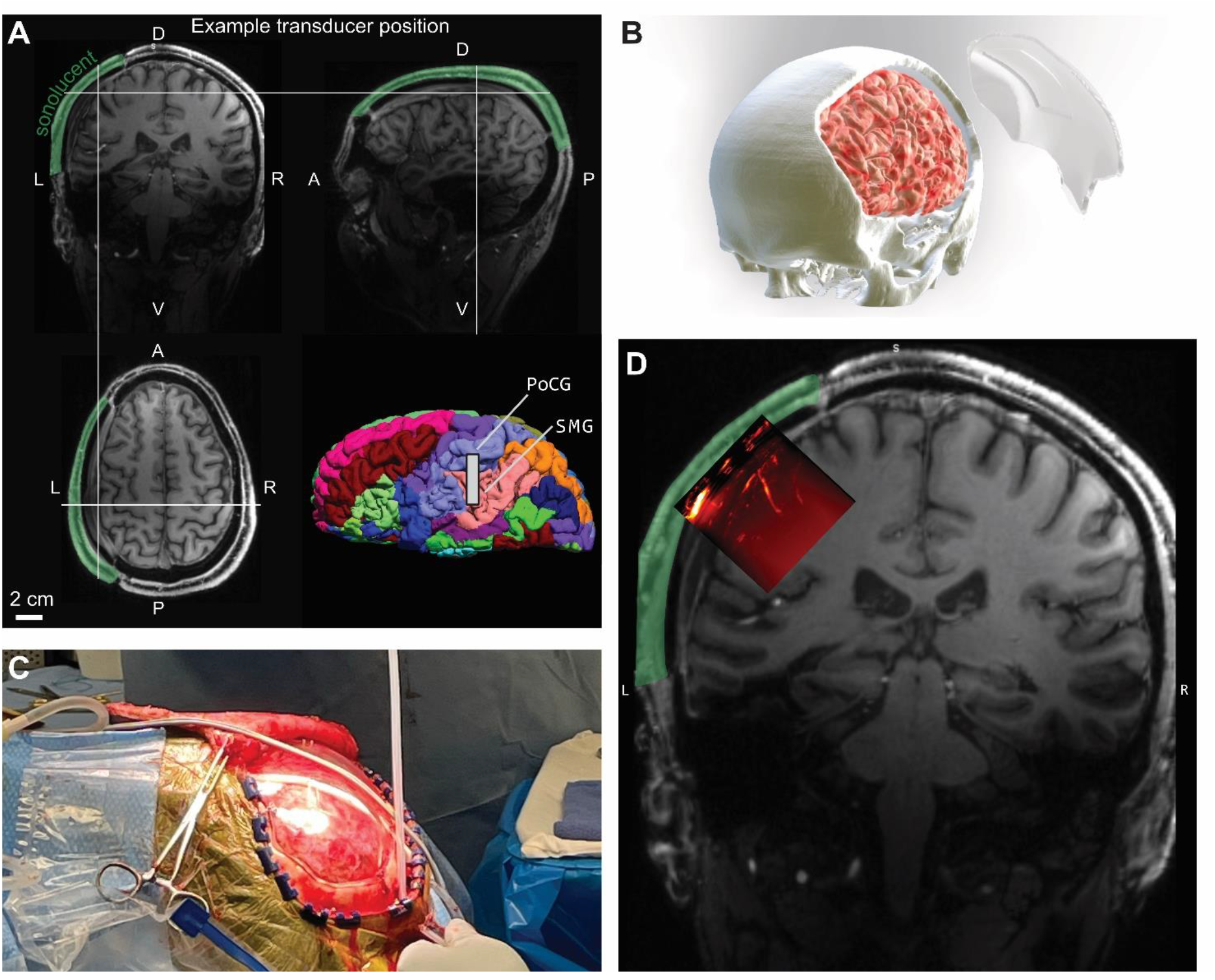
Participant J was reconstructed with a custom-designed permanent acoustic window. (**A**) MRI scan of Participant J after reconstruction. The white crosses indicate the middle of the transducer during the example fUSI session. Green shading - sonolucent portion of head, including scalp, CCI, and meninges above brain. White bar in bottom right brain: estimated position of the transducer. SMG: Supramarginal gyrus; PoCG: Postcentral gyrus (**B**) 4mm thick sonolucent cranial implant with the 2 mm thick parallelogram-shaped “window” placed above the primary motor, primary somatosensory, and posterior parietal cortex (**C**) Reconstruction surgery of Participant J with the PMMA CCI (D) Co-registration of the fUSI imaging plane with an anatomical MR image.

### Acoustic window allows fUSI activity recording in fully reconstructed human participant

We imaged Participant J’s brain through his acoustic window following his skull reconstruction (**Fig. 5C**). We first located the boundaries of the thinned window using real-time anatomical B-mode ultrasound imaging. Once we located the boundaries of the “window”, we used a custom-designed cap to stably position the ultrasound transducer above the middle of the acoustic 2-mm window. We immediately observed the cortical vasculature, including vessels following the curves of sulcal folds and smaller vessels irrigating the adjacent cortex (**Fig. 5D**).

Based on a prior fUSI recording session and the location of the thinned window, we estimated that the transducer was positioned above the left primary somatosensory cortex (S1) and supramarginal gyrus (SMG), with S1 playing a role in processing somatic sensory signals from the body^15, 16^ and the SMG playing a role in grasping and tool use^17–23^. Thus, in an attempt to detect functional brain signals, we instructed Participant J to perform two visuomotor tasks while sitting in a comfortable chair with a screen in front of him (**Fig. 6A**). In the first task, we used a block design with 100-second “rest” blocks and 50-second “task” blocks. During the rest blocks, we instructed the participant to close his eyes and relax. During the task blocks, the participant used a video game controller joystick. He was instructed to complete “connect-the-dots” puzzles on the computer monitor (**Fig. 6B**). He used his right thumb to control the game controller’s thumbstick (cursor location) and his left index finger to control the left shoulder button (mouse click). We repeated the same tasks across multiple runs (N=3). Finally, we concatenated the data from two runs and used a GLM analysis to identify voxels with functional activation. The GLM revealed several regions that were task-modulated (**Fig. 6C**,**D**). The activity within these regions displayed positive modulation by the task, i.e., increased activity during the drawing blocks and decreased activity during the rest blocks (ROI 2, **Fig. 6F**). For example, ROI 2 had an average of 3.68% difference between the drawing and rest blocks (p value <10^-10^, two-sided t-test). Outside of these activated regions (i.e., ROI 1, **Fig. 6F**), the signal remained stable throughout the run with no significant increase nor decrease during the task periods. For example, ROI 1 had an average difference of -0.034% between the drawing and rest blocks, (p value = 0.67, two-sided t-test).

**Figure 6.**
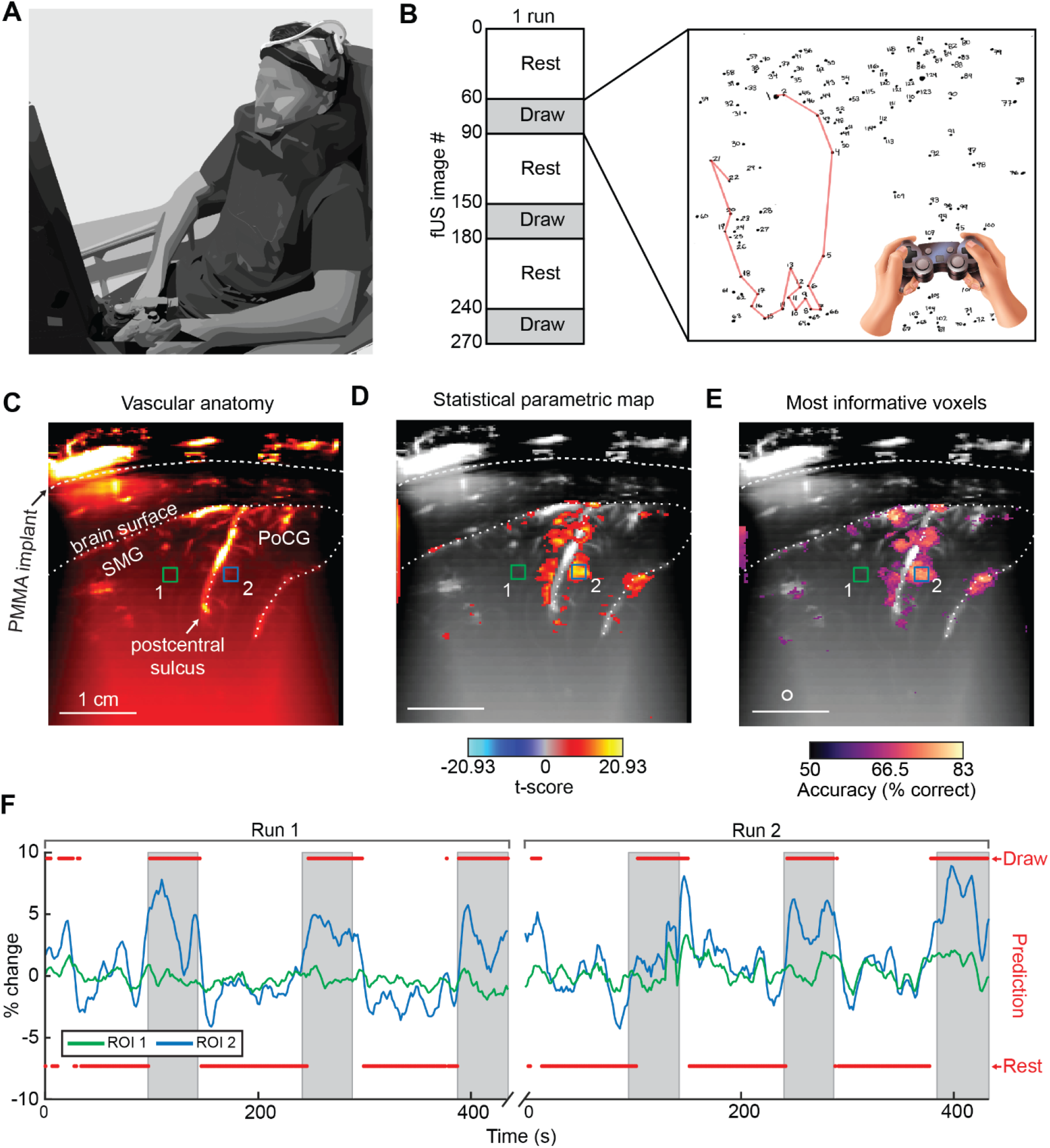
Permanent acoustic window allows non-invasive fUSI imaging and decoding during a gaming task. (**A**) Example setup of Participant J playing connect-the-dots with a joystick during fUSI recording (**B**) Connect-the-dot task. In the rest blocks, the participant relaxed and tried to keep his mind clear. In the task blocks, the participant used a game controller to draw lines in a “connect-the-dots” task (**C**) Vascular anatomy of the imaging plane. Dashed lines highlight specific anatomic features, including PMMA implant surface, brain surface, and sulcal vessels. SMG: Supramarginal gyrus; PoCG: Postcentral gyrus. Colored boxes show ROIs used in part F (**D**) Task-modulated areas across two concatenated runs. T-score statistical parametric map, values shown for voxels where p(corrected) < 10-10 (E) Searchlight analysis. Top 5 % of voxels with the highest decoding accuracy. White circle – 600 μm searchlight radius. Masked voxels correspond to threshold of p(corrected) < 2.8×10-4 (F) Mean scaled fUSI signal from ROIs. White regions are rest blocks; grey regions are task blocks. Red circles show prediction from linear decoder.

As a very first step towards human BMI applications, we tested the ability to decode task state (rest vs. connect-the-dot) from single trials of the fUSI data using a linear decoder. We successfully decoded the task state with 84.7 % accuracy (p<10^-15^, 1-sided binomial test). To better understand which voxels in the image contained the most information discriminating the task blocks, we performed a searchlight analysis with a 600 μm radius (**Fig. 6E**). This analysis showed that the 5% most informative voxels were distributed across the image and closely matched the results of the statistical parametric map from the GLM. When we examined the decoder accuracy across our example session, our linear decoder predicted both the draw and rest blocks with similarly high accuracy (**Fig. 6F**), with most of the errors occurring at the transitions between the two task states. This effect is likely due in part to the latency between the neural activity and resulting hemodynamic response^24, 25^.

In the second task, we asked Participant J to play guitar while we recorded fUSI data (**Fig.7A**,**B**). During the rest blocks (100-second), we instructed him to minimize finger/hand movements, close his eyes, and relax. During the task blocks (50-second), the participant played improvised or memorized music on a guitar with his right-hand strumming and his left fingers moving on the fretboard. We identified several regions that were task-activated, including several that were similar in location to those activated by the connect-the-dots task (**Fig. 7C**,**D**).

**Figure 7.**
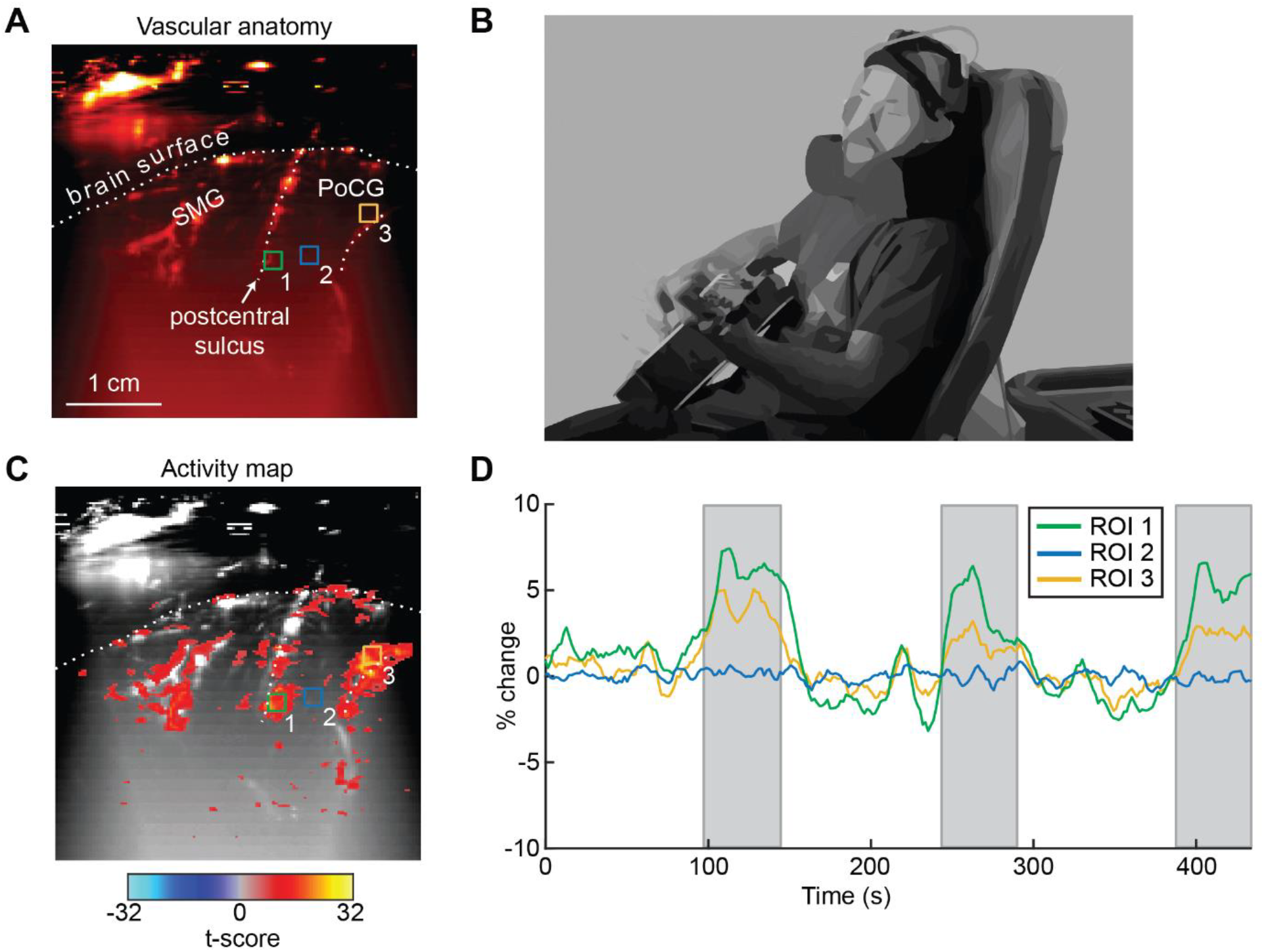
Permanent acoustic window allows fUSI detection of guitar playing. (A) Vascular anatomy of the imaging plane. SMG: Supramarginal gyrus; PoCG: Postcentral gyrus (B) Participant J playing guitar during fUSI recording. Colored boxes show ROIs used in part D (C) Guitar playing-modulated areas. T-score statistical parametric map thresholded at p(corrected) < 10-10 (D) Mean scaled fUSI signal from ROIs. White regions are rest blocks; grey regions are task blocks.

## DISCUSSION

fUSI presents a host of benefits (e.g., increased sensitivity, resolution, and portability) relative to more established techniques such as fMRI. However, fUSI cannot penetrate human skull bone while maintaining sufficient sensitivity. In this study, we established, for the first time, the feasibility of awake human fUSI imaging in a non-surgical setting through a polymeric acoustic window. Before testing this approach in humans, we characterized the acoustic performance of the reconstruction material in vitro (**Fig. 2**) and in vivo (rodent) (**Fig. 3**) settings. This allowed us to determine the feasibility of non-invasive imaging through a CCI and design an appropriate acoustic window. We subsequently acquired, for the first time, functional activity of the brain using fUSI in an awake, behaving adult participant completely noninvasively, outside of a surgery environment (**Fig. 6, Fig. 7**). We additionally demonstrate, for the first time, decoding of human brain states associated with different tasks. Notably, our success in using fUSI to decode brain states (**Fig. 6**) serves as a crucial initial precursor to ultrasonic brain-machine interfaces in humans. Furthermore, our overall approach opens numerous potential applications in both research and clinical use. The following paragraphs expand on these possibilities and their potential impact.

### Diagnostics & monitoring after skull reconstruction (clinical use)

It is currently difficult and expensive to monitor anatomical and functional brain recovery following a cranioplasty. Behavioral assessments, such as Cognitive Status Examination, Mini-Mental State Examination, or Functional Independence Measure are commonly used to assess neuropsychological recovery following traumatic brain injuries^26–28^ but cannot identify specific sites of damage or track recovery at these anatomical locations. Less commonly, CT and/or MRI are used to assess anatomical and functional recovery^29^. However, these methods possess low sensitivity/specificity for assessing brain recovery, are expensive (CT + MRI), and can add risk to the patient (CT). In the future, fUSI and CCIs with acoustic windows may enable routine monitoring during the postoperative period for both anatomical and functional recovery. In addition to generalized post-operative monitoring, some TBI patients will develop specific pathologies that would benefit from increased monitoring frequency. For example, Syndrome of the Trephined (SoT) is an indication where patients develop neurological symptoms such as headaches, dizziness, and cognitive impairments due to altered cerebrospinal fluid dynamics and changes in intracranial pressure following a large craniectomy^30^. Recording from these patients with TBI sequelae or SoT may provide novel insight into the pathophysiology of their disease processes and subsequent recovery.

### Neuroscience and brain-machine interfaces (research use)

One of the most significant bottlenecks to human neuroscience research and the development of less invasive BMI is the limited access to human patients for obtaining neural activity data. The ability to measure fUSI signals from fully reconstructed, ambulatory adult humans has the potential to address this challenge, opening opportunities for advancements in these research areas. Approximately 1.7 million people suffer from a severe traumatic brain injury each year in the United States^31^. If only a small fraction of these patients receives a cranial implant with an acoustic window as part of their standard of care, it would provide a major opportunity to measure mesoscopic neural activity with excellent spatiotemporal resolution and high sensitivity in humans. In those patients with minimal long-term neurological damage, it will also enable new investigations into advanced neuroimaging techniques and BMI. As we demonstrated in this paper, fUSI possesses high sensitivity even through the acoustic window. Not only could we identify task-modulated areas by averaging across all task blocks and using a GLM (**Fig. 6D**), but we could also use a linear decoder to robustly decode the current task block using single fUSI images (**Fig. 6E**,**F**).

## CONCLUSION

Taken together, our results suggest that acoustic windows for fUSI could bridge the gap between existing high-precision but highly-invasive and non-invasive but lower-precision technologies for neural recording. The large field-of-view (38 mm x 50 mm), high spatial precision (200 μm) and sensitivity (single-trial decoding) demonstrated by this technology provides unprecedented access to brain activity in fully reconstructed adult humans. This access has the potential to directly benefit brain injury patients and open new doors to neuroscience discoveries and the development of improved treatments and BMIs.

## MATERIALS AND METHODS

### General

All analysis was completed in MATLAB 2021a. Implant materials

PMMA – 1, 2, 3, 4 mm. Provided by Longeviti.

Titanium mesh – Pure Titanium, 0.6 mm thick, honeycomb patterns alternating between small circle (1.5 mm diameter) and big circle (3 mm diameter); KLS Martin

### Functional ultrasound imaging: (fUSI) mode

Functional ultrasound imaging (fUSI) visualizes neural activity by mapping local changes in cerebral blood volume (CBV). CBV variations are tightly linked to neuronal activity through the neurovascular coupling^32^ and are evaluated by calculated power Doppler variations in the brain^3^. fUSI used an ultrasonic probe centered at 7.5 MHz (Bandwidth > 60%, 128 elements, 0.300 mm pitch, Vermon, Tours, France) connected to a Verasonics Vantage ultrasound system (Verasonics Inc., Redmond, WA, USA) controlled by custom MATLAB (MathWorks, USA) B-mode and fUSI acquisition scripts. Each power Doppler image was obtained from the accumulation of 300 compounded frames acquired at 400 Hz frame rate. Each compounded frame was created using 2 accumulations of 5 tilted plane waves (-6°,-3°, 0°, 3°, 6°). We used a pulse repetition frequency (PRF) of 4000 Hz. fUSI images were repeated every 1.65 seconds. Each block of 300 images was processed using a SVD clutter filter^33^ to separate tissue signal from blood signal to obtain a final power Doppler image exhibiting artificial (for in vitro experiment) or cerebral blood volume (CBV) in the whole imaging plane (**Fig. 1E**).

### In vitro tissue anatomical and doppler phantoms

We routed 280 μm inner diameter polyethylene tubing through a hollow, box-shaped, 3D-printed, nylon cast at three lateral positions and five axial positions (15 grid points, total). We then poured a liquid gelatin phantom with graphite added to mimic the scattering effects of biological soft tissue. Once the phantom cast had set/solidified, we flowed a red blood cell phantom liquid (CAE Blue Phantom™ Doppler Fluid) through the tubing using a peristaltic pump and a long recirculating route with a low pass filter to create a smooth flow at velocities of approximately 0.1 mL/min.

### In vivo functional ultrasound imaging comparative study in rat

Four Long-Evans male rats were used in this study (15-20 weeks old, 500 g–650 g, Caltech protocol number: IA22-1729). During the surgery and the subsequent imaging session, the animals were anesthetized using an initial intraperitoneal injection of xylazine (10 mg/kg) and ketamine (Imalgene, 80 mg/kg). The scalp of the animals was removed, and the skull was clean with saline. A craniectomy was performed to remove 0.5mm × 1 cm of the skull by drilling (Foredom) at low speed using a micro drill steel burr (Burr number 19007-07, Fine Science Tools). We took care to avoid damage to the dura and prevent brain inflammation. After surgery, the surface of the brain was rinsed with sterile saline and ultrasound coupling gel was placed in the window. The linear ultrasound transducer was positioned directly above the cranial window and a fUSI scan was performed. We then placed the 1 mm-2 mm-, 3 mm-thick PMMA materials or the titanium mesh, above the brain, and repeated the fUSI acquisition.

To quantitatively characterize the fUSI sensitivity through the different PMMA thicknesses, we calculated blood vessels SNR in the cortex and in deeper thalamic regions from the same animal with different implants. **Figure S1.a** shows the two regions of interest (ROIs) selected for each implant condition (i. Cortex, ii. Deeper regions). For each horizontal line of these ROIs, the lateral intensity was plotted, and local maxima (blood vessels) and minima were identified (**Fig. S1.b-c**). SNR was then calculated as : 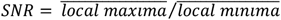.

fUSI with visual stimuli was performed in one animal. Visual stimuli were delivered using a blue LED (450 nm wavelength) positioned at 5 cm in front of the head of the rat. Stimulation runs consisted of periodic flickering of the blue LED (flickering rate: 5 Hz) using the following parameters: 50 s dark, followed by 16.5 s of light flickering repeated three times for a total duration of 180 s. At this distance, the light luminance was of 14 lux when the light was on and ∼0.01 lux when the light was off.

### fUSI data processing

For the rodent and human in vivo experiments, we used a General Linear Model (GLM) to find which voxels were significantly modulated by the visual task. To perform this GLM, we first preprocessed the fUSI data with rigid body motion correction^34^, followed by spatial smoothing (2D Gaussian with sigma=1 (FWHM = 471 μm), followed by a voxelwise moving average temporal filter (rat: 2-timepoints; human: 5-timepoints). We then scaled the fUSI signal by its voxelwise mean so that all the runs and voxels had a similar signal range^35^. To generate the GLM regressor for the visual task, we convolved the block task design with a single Gamma hemodynamic response function (HRF)^36^. For the rodent experiments, the HRF time constant was (τ)=0.7, time delay (δ)=1 s, and phase delay (n)=3 s. For the human experiments, the values were τ=0.7, δ=3 s, n=3 s. We next fit the GLM using the convolved regressor and the scaled fUSI signal from each voxel. We determined statistical significance of the beta coefficients for each voxel using a 2-sided t-test with False Discovery Rate (FDR) correction (p (corrected) < 10^-5^).

### Human participant

We recruited and consented a 35-year-old male participant (J) with a traumatic brain injury to participate in a research study examining the ability to record functional ultrasound signals through a custom cranial implant. All procedures were approved by the Institutional Review Boards (IRB) of the University of Southern California (USC), California Institute of Technology (Caltech), and Rancho Los Amigos National Rehabilitation Hospital (RLA). Caltech reference number IR19-0902. All fUSI study sessions took place at Caltech. All CT and MRI scans occurred at the Keck Hospital of USC. Decompressive hemicraniectomy and reconstruction procedure Patient J underwent a decompressive hemicraniectomy following a severe traumatic brain injury on April 9, 2019. The approximate size of the craniectomy was 16 cm in anterior-posterior by 10 cm dorsal-ventral (**Fig. 4B**). A 700-μm isotropic anatomical MRI was acquired shortly after the hemicraniectomy. Patient J underwent a left cranioplasty using the Longeviti ClearFit custom skull implant on September 22, 2021. The surgery was performed at Rancho Los Amigos National Rehabilitation Center. The surgery was completed in the standard fashion. Briefly, after general anesthesia was induced, the left side of the head was prepped and draped. The prior hemicraniectomy incision was opened and the scalp was dissected from the dura and the edges of the skull defect was identified circumferentially. An epidural surgical drain was placed, and the cranioplasty implant was secured to the skull using titanium microplates and screws before the wound was closed in multiple layers. The surgical drain was placed to minimize the accumulation of epidural fluid and was removed after three days.

### Skull implant design

The PMMA skull implant (Longeviti ClearFit) was designed to fit the hemicraniectomy and match the geometry of the right side of the intact skull. The implant was 4-mm thick to match the patient’s nominal bone thickness except for a 34 mm x 50 mm parallelogram-shaped “window” of 2 mm thick PMMA positioned over the area of the brain known to be active during finger tapping, based on the results of a functional MRI experiment.

### Human fMRI task

Participant J participated in a fMRI scan during which he performed a finger tapping task with a block design of 30 s rest followed by 30 s sequential finger tapping with his right hand. These blocks were repeated 7 times for a total scan duration of 8 min. Instructions for start and end of finger tapping epochs were delivered with auditory commands delivered through MR compatible headphones. The fMRI acquisition was done on a 7T Siemens Magnetom Terra system with a 32-channel receive 1Tx head coil with a multi-band gradient echo planar imaging (GE-EPI) T2*-weighted sequence with 1 mm^3^ isotropic resolution, 192 mm x 192 mm FOV, 92 axial slices, TR/TE 3000 ms/22 ms, 160 volumes, FA 80 deg, A-P phase encoding direction, iPAT=3 and SMS=2. An anatomical scan was also acquired using a T1-weighted MPRAGE sequence with 0.7 mm^3^ isotropic resolution, 224 mm x 224 mm FOV, 240 sagittal slices, TR/TE 2200 ms/2.95 ms, FA 7 deg. Statistical analysis of fMRI data was performed with a General-Linear Model using Statistical Parametric Mapping (SPM12). Preprocessing included motion-realignment, linear drift removal and co-registration of fMRI data to high resolution anatomical scan.

### Human fUSI tasks

Participant J was seated in a reclining chair with a 27-inch fronto-parallel screen (Acer XB271HU) positioned 70 cm in front of him. Participant J controlled the behavioral task using a Logitech F310 Gamepad. We used Gopher (https://github.com/Tylemagne/Gopher360) to enable control of the computer with the Logitech Gamepad. Right thumbstick controlled the position of the computer cursor while the left shoulder button functioned as the left mouse button. We used a block design for the drawing task with 100-second rest blocks followed by 50 seconds of drawing with the Gamepad. We verbally instructed the participant for each rest or task block. The participant was instructed to complete one of multiple “Connect-The-Dots” drawings (**Fig. 6B**). When the participant finished one of the drawings, we presented a new drawing for him to complete. For the rest blocks, we instructed the participant to close his eyes and try to keep his mind relaxed. We acquired fUSI data at 0.6 Hz (1.65 sec/frame).

For the guitar playing task, we used an identical block design with 60-frame rest blocks followed by 30-frame task blocks. In the task blocks, the participant used his left hand to form chords on the fretboard and his right hand to strum the strings.

### Task decoding

To decode whether a given timepoint was in a “task” or “rest” block, we used principal component analysis (PCA) for dimensionality reduction and linear discriminant analysis (LDA) for classification. We first labeled each motion-corrected fUSI timepoint (“sample”) as “rest” or “task”. We then balanced the dataset to have an equal number of “rest” and “task” timepoints. We then split the dataset at the level of block pairs (1 block pair = rest+task) to avoid training the classifier on time points immediately adjacent to the test time points. This helps ensure that the model would generalize and that our model was not memorizing local patterns for each block pair. We then applied a 2D Gaussian smoothing filter (sigma=1) to each sample in the train and test sets. We z-scored the train set across time for each voxel. We then trained and validated the PCA+LDA classifier using a block-wise leave-one-out cross-validator; i.e., we trained on 5 blocks and then tested on the held-out block pair’s timepoints. For the PCA, we kept 95 % of the variance. To generate the example session decoding, we trained on 5 blocks with balanced samples of “rest” and “draw” and then tested on the unbalanced final block (60 fUSI frames of rest data and 30 fUSI frames of draw task).

### Searchlight analysis

We defined a circular region of interest (ROI) and, using only the pixels within the ROI, we performed the task decoding analysis. We assigned that ROI’s percent correct metric to the center voxel. We then repeated this across the entire image, such that each image pixel is the center of one ROI. To visualize the results, we overlaid the percent correct metric onto a vascular map and kept the 5% most significant voxels. We only ran the searchlight analysis on brain voxels, i.e., we ignored all voxels above the brain surface.

## ACKNOWLEDGMENTS

We would like to thank Kelsie Pejsa for administrative assistance and participant planning. We would also like to thank Professor Mickael Tanter at INSERM (Paris, France) for his feedback and support throughout the research process. Finally, we would like to thank Participant J for his willing participation.

## FUNDING

NIH (R01NS123663): RAA, MGS

The T&C Chen Brain-Machine Interface Center: RAA, MGS The Boswell Foundation: RAA

NEI F30 (NEI F30 EY032799): WSG

Josephine de Karman Fellowship: WSG

UCLA-Caltech MSTP (NIGMS T32 GM008042): WSG

Della Martin Postdoctoral Fellowship: SLN

Human Frontier Science Program Cross-Disciplinary Fellowship (LT000217/2020-C) : CR

USC Neurorestoration Center : CL

## AUTHOR CONTRIBUTIONS

Conceptualization: CR, SLN, WSG, CL, RAA, MGS fUSI sequence development: CR, SLN

Doppler phantom design: SLN In vitro experiments: CR, SLN

Rodent experiments: CR

Study participant recruitment : CL Human fUSI recording: CR, SLN, WSG

Craniectomy and cranioplasty surgeries: CL, JJR Structural and functional MR imaging: KJ

MR imaging analysis: KJ

Ultrasound data processing and analysis: WSG, SLN, CR Supervision of the research : MGS, RAA, CL,VC Writing – original draft : CR, WSG, SLN, MGS, RAA

Writing – review & editing: CR, WSG, SLN, MGS, RAA, CL,VC

## DATA AND MATERIALS AVAILABILITY

Key data used in this paper will be stored on DANDI (https://www.dandiarchive.org/), CaltechDATA (https://data.caltech.edu/), or a similar data repository. DOI accession number will be generated upon acceptance of paper. Code used to generate key figures and results will be posted to a publicly accessible GitHub repository and an archived version will be stored on Zenodo or similar archivable code repository. DOI accession number will be generated upon acceptance of paper.

## REFERENCES

1. E. Macé, G. Montaldo, I. Cohen, M. Baulac, M. Fink, M. Tanter, Functional ultrasound imaging of the brain. Nat Methods 8, 662–664 (2011).

2. T. Deffieux, C. Demene, M. Pernot, M. Tanter, Functional ultrasound neuroimaging: a review of the preclinical and clinical state of the art. Current Opinion in Neurobiology 50, 128–135 (2018).

3. C. Rabut, S. Yoo, R. C. Hurt, Z. Jin, H. Li, H. Guo, B. Ling, M. G. Shapiro, Ultrasound Technologies for Imaging and Modulating Neural Activity. Neuron 108, 93–110 (2020).

4. S. L. Norman, D. Maresca, V. N. Christopoulos, W. S. Griggs, C. Demene, M. Tanter, M. G. Shapiro, R. A. Andersen, Single Trial Decoding of Movement Intentions Using Functional Ultrasound Neuroimaging. bioRxiv, 2020.05.12.086132 (2020).

5. W. S. Griggs, S. L. Norman, T. Deffieux, F. Segura, B.-F. Osmanski, G. Chau, V. Christopoulos, C. Liu, M. Tanter, M. G. Shapiro, R. A. Andersen, Decoding Motor Plans Using a Closed-Loop Ultrasonic Brain-Machine Interface, 2022.11.10.515371 (2022).

6. C. Brunner, M. Grillet, A. Urban, B. Roska, G. Montaldo, E. Macé, Whole-brain functional ultrasound imaging in awake head-fixed mice. Nat Protoc 16, 3547–3571 (2021).

7. M. Imbault, D. Chauvet, J.-L. Gennisson, L. Capelle, M. Tanter, Intraoperative Functional Ultrasound Imaging of Human Brain Activity. Sci Rep 7, 7304 (2017).

8. S. Soloukey, A. J. P. E. Vincent, D. D. Satoer, F. Mastik, M. Smits, C. M. F. Dirven, C. Strydis, J. G. Bosch, A. F. W. van der Steen, C. I. De Zeeuw, S. K. E. Koekkoek, P. Kruizinga, Functional Ultrasound (fUS) During Awake Brain Surgery: The Clinical Potential of Intra-Operative Functional and Vascular Brain Mapping. Front. Neurosci. 13 (2020), doi:10.3389/fnins.2019.01384.

9. C. Demene, J. Baranger, M. Bernal, C. Delanoe, S. Auvin, V. Biran, M. Alison, J. Mairesse, E. Harribaud, M. Pernot, M. Tanter, O. Baud, Functional ultrasound imaging of brain activity in human newborns. Sci. Transl. Med. 9, eaah6756 (2017).

10. H. Alvis-Miranda, S. M. Castellar-Leones, L. R. Moscote-Salazar, Decompressive Craniectomy and Traumatic Brain Injury: A Review. Bull Emerg Trauma 1, 60–68 (2013).

11. E. Güresir, P. Schuss, H. Vatter, A. Raabe, V. Seifert, J. Beck, Decompressive craniectomy in subarachnoid hemorrhage. Neurosurgical Focus 26, E4 (2009).

12. L.-P. Pallesen, K. Barlinn, V. Puetz, Role of Decompressive Craniectomy in Ischemic Stroke. Frontiers in Neurology 9 (2019) (available at https://www.frontiersin.org/articles/10.3389/fneur.2018.01119).

13. C. Iaccarino, A. G. Kolias, L.-G. Roumy, K. Fountas, A. O. Adeleye, Cranioplasty Following Decompressive Craniectomy. Front Neurol 10, 1357 (2020).

14. T. Shay, K.-A. Mitchell, M. Belzberg, I. Zelko, S. Mahapatra, J. Qian, L. Mendoza, J. Huang, H. Brem, C. Gordon, Translucent Customized Cranial Implants Made of Clear Polymethylmethacrylate: An Early Outcome Analysis of 55 Consecutive Cranioplasty Cases. Ann Plast Surg 85, e27–e36 (2020).

15. B. P. Delhaye, K. H. Long, S. J. Bensmaia, in Comprehensive Physiology, R. Terjung, Ed. (Wiley, 2018), pp. 1575–1602.

16. W. Penfield, E. Boldrey, Somatic motor and sensory representation in the cerebral cortex of man as studied by electrical stimulation. Brain 60, 389–443 (1937).

17. S. K. Wandelt, S. Kellis, D. A. Bjånes, K. Pejsa, B. Lee, C. Liu, R. A. Andersen, Decoding grasp and speech signals from the cortical grasp circuit in a tetraplegic human. Neuron 110, 1777–1787.e3 (2022).

18. G. A. Orban, F. Caruana, The neural basis of human tool use. Frontiers in Psychology 5 (2014) (available at https://www.frontiersin.org/articles/10.3389/fpsyg.2014.00310).

19. J. P. Gallivan, D. A. McLean, K. F. Valyear, J. C. Culham, D. Angelaki, Ed. Decoding the neural mechanisms of human tool use. eLife 2, e00425 (2013).

20. T. McDowell, N. P. Holmes, A. Sunderland, M. Schürmann, TMS over the supramarginal gyrus delays selection of appropriate grasp orientation during reaching and grasping tools for use. Cortex: A Journal Devoted to the Study of the Nervous System and Behavior 103, 117–129 (2018).

21. M. Buchwald, Ł. Przybylski, G. Króliczak, Decoding Brain States for Planning Functional Grasps of Tools: A Functional Magnetic Resonance Imaging Multivoxel Pattern Analysis Study. Journal of the International Neuropsychological Society 24, 1013–1025 (2018).

22. F. E. Garcea, L. J. Buxbaum, Gesturing tool use and tool transport actions modulates inferior parietal functional connectivity with the dorsal and ventral object processing pathways. Hum Brain Mapp 40, 2867–2883 (2019).

23. S. K. Wandelt, D. A. Bjånes, K. Pejsa, B. Lee, C. Liu, R. A. Andersen, Online internal speech decoding from single neurons in a human participant, 2022.11.02.22281775 (2022).

24. J. Claron, M. Provansal, Q. Salardaine, P. Tissier, A. Dizeux, T. Deffieux, S. Picaud, M. Tanter, F. Arcizet, P. Pouget, Co-variations of cerebral blood volume and single neurons discharge during resting state and visual cognitive tasks in non-human primates. Cell Reports 42, 112369 (2023).

25. A. O. Nunez-Elizalde, M. Krumin, C. B. Reddy, G. Montaldo, A. Urban, K. D. Harris, M. Carandini, Neural correlates of blood flow measured by ultrasound. Neuron 110, 1631–1640.e4 (2022).

26. N. A. Nabors, S. R. Millis, M. Rosenthal, Use of the Neurobehavioral Cognitive Status Examination (Cognistat) in Traumatic Brain Injury. The Journal of Head Trauma Rehabilitation 12, 79 (1997).

27. E. de Guise, N. Gosselin, J. LeBlanc, M.-C. Champoux, C. Couturier, J. Lamoureux, J. Dagher, J. Marcoux, M. Maleki, M. Feyz, Clock Drawing and Mini-Mental State Examination in Patients with Traumatic Brain Injury. Applied Neuropsychology 18, 179–190 (2011).

28. K. Smith-Knapp, J. D. Corrigan, J. A. Arnett, Predicting functional independence from neuropsychological tests following traumatic brain injury. Brain Injury 10, 651–662 (1996).

29. R. S. Scheibel, Functional Magnetic Resonance Imaging of Cognitive Control following Traumatic Brain Injury. Front Neurol 8, 352 (2017).

30. K. Ashayeri, E. M. Jackson, J. Huang, H. Brem, C. R. Gordon, Syndrome of the Trephined: A Systematic Review. Neurosurgery 79, 525–534 (2016).

31. A. Georges, J. M. Das, Traumatic Brain Injury (StatPearls Publishing, 2022; https://www.ncbi.nlm.nih.gov/books/NBK459300/).

32. C. Iadecola, The Neurovascular Unit Coming of Age: A Journey through Neurovascular Coupling in Health and Disease. Neuron 96, 17–42 (2017).

33. C. Demené, T. Deffieux, M. Pernot, B.-F. Osmanski, V. Biran, J.-L. Gennisson, L.-A. Sieu, A. Bergel, S. Franqui, J.-M. Correas, I. Cohen, O. Baud, M. Tanter, Spatiotemporal Clutter Filtering of Ultrafast Ultrasound Data Highly Increases Doppler and fUltrasound Sensitivity. IEEE Transactions on Medical Imaging 34, 2271–2285 (2015).

34. E. A. Pnevmatikakis, A. Giovannucci, NoRMCorre: An online algorithm for piecewise rigid motion correction of calcium imaging data. Journal of Neuroscience Methods 291, 83–94 (2017).

35. G. Chen, P. A. Taylor, R. W. Cox, Is the statistic value all we should care about in neuroimaging? NeuroImage 147, 952–959 (2017).

36. G. M. Boynton, S. A. Engel, G. H. Glover, D. J. Heeger, Linear Systems Analysis of Functional Magnetic Resonance Imaging in Human V1. J. Neurosci. 16, 4207–4221 (1996).

