## Supplementary Information for "A window to the brain: ultrasound imaging of human neural activity through a permanent acoustic window"

[1]Division of Chemistry and Chemical Engineering, California Institute of Technology; Pasadena, CA 91125, USA
[2] Division of Biology and Biological Engineering, California Institute of Technology; Pasadena, CA 91125, USA
[3] USC Neurorestoration Center and the Departments of Neurosurgery and Neurology, University of Southern California; Los Angeles, CA 90033, USA
[4] Stevens Neuroimaging and Informatics Institute, University of Southern California; Los Angeles, CA 90033, USA
[5] Department of Bioengineering, University of California Riverside; Riverside, CA 92521, USA
[6] The T&C Chen Brain-machine Interface Center, California Institute of Technology; Pasadena, CA 91125, USA
[7] Howard Hughes Medical Institute; Pasadena, CA 91125, USA
[†] These authors contributed equally

### SUPPLEMENTAL DATA

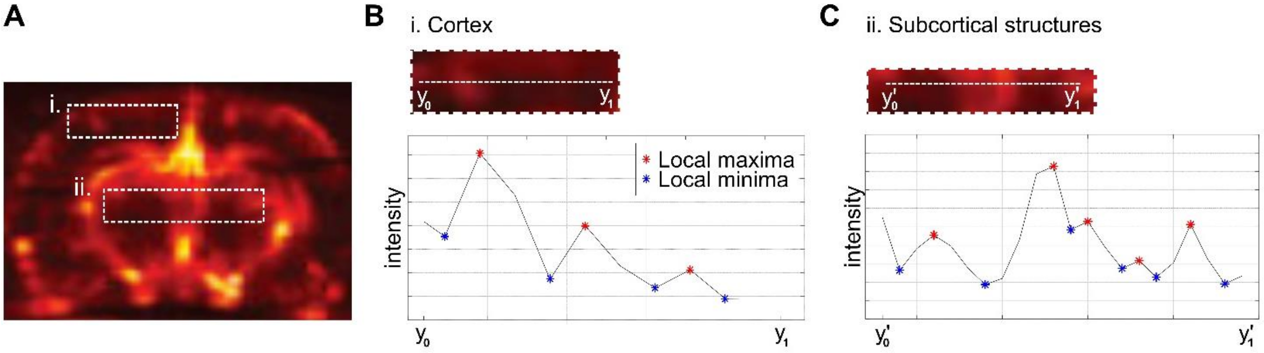


**Fig. S1: SNR calculation for in vivo characterization of PMMA as a sonolucent implant for fUSI** (**A**) For each condition (implant material, thickness, etc.) two regions are analyzed: the cortex (i.) and the deeper regions (ii.) (**B,C**) For both regions, the local maxima (red stars) and local minima (blue stars) are calculated along each depth to determine the SNR.
